# Orientation processing by synaptic integration across first-order tactile neurons

**DOI:** 10.1101/396705

**Authors:** Etay Hay, J Andrew Pruszynski

**Author notes:** Correspondence: Dr. Etay Hay, Krembil Centre for Neuroinformatics, Centre for Addiction and Mental Health, 250 College St, Toronto, Ontario, M5T 1R8.

## Abstract

Our ability to manipulate objects relies on tactile inputs from first-order tactile neurons that innervate the glabrous skin of the hand. The distal axon of these neurons branches in the skin and innervates many mechanoreceptors, yielding spatially-complex receptive fields. Here we show that synaptic integration across the complex signals from the first-order neuronal population could underlie human ability to accurately (< 3°) and rapidly process the orientation of edges moving across the fingertip. We first derive spiking models of human first-order tactile neurons that fit and predict responses to moving edges with high accuracy. We then use the model neurons in simulating the peripheral neuronal population that innervates a fingertip. We use machine learning to train classifiers performing synaptic integration across the neuronal population activity, and show that synaptic integration across first-order neurons can process edge orientations with high acuity and speed. In particular, our models suggest that integration of fast-decaying (AMPA-like) synaptic inputs within short timescales is critical for discriminating fine orientations, whereas integration of slow-decaying (NMDA-like) synaptic inputs refine discrimination and maintain robustness over longer timescales. Taken together, our results provide new insight into the computations occurring in the earliest stages of the human tactile processing pathway and how they may be critical for supporting hand function.

**Author Summary:** Our ability to manipulate objects relies on tactile inputs signaled by first-order neurons that innervate mechanoreceptors in the skin of the hand and have spatially-complex receptive fields. Here we show how synaptic integration across the rich inputs from first-order neurons can rapidly and accurately process the orientation of edges moving across the fingertip. We derive spiking models of human first-order tactile neurons, then use the models to simulate the peripheral neuronal population that innervates a fingertip. We use machine learning to train classifiers performing synaptic integration across the neuronal population activity, and show that synaptic integration across first-order neurons could underlie human ability to process edge orientations with high acuity and speed.

## Introduction

Tactile input from the hands is important for many behaviors, ranging from daily motor tasks like buttoning a shirt to complex skills like knitting a sock [1]. This tactile information is conveyed by four types of first-order tactile neurons that innervate distinct mechanoreceptive end organs in the glabrous skin of the hand [2]. Of particular note are fast-adapting type 1 neurons (FA-1) innervating Meissner corpuscles and slow-adapting type 1 neurons (SA-1) innervating Merkel discs, which are particularly important for extracting fine spatial details of touched objects [3]. A fundamental feature of FA-1 and SA-1 neurons is that their axon branches in the skin and innervates many mechanoreceptors, ~30 on average [4], and thus these neurons have spatially-complex receptive fields with many highly-sensitive zones [3,5–7]. We have recently proposed that this innervation pattern may constitute a peripheral neural mechanism for extracting geometric features of touched objects [6].

Here we examine how the geometric features of touched objects are encoded by the population of first-order tactile neurons and the synaptic readout of their activity. Our approach adds to previous studies [8–12] in several ways. First, our models are based on first-order tactile neurons that have spatially complex receptive fields, and involve spiking and other physiological details that were absent in previous simple models of tactile receptive field responsivity [6]. Second, our models are constrained by neural recordings in humans, whereas previous models used data from monkeys [9,10]. Third, previous models of the peripheral neuronal population innervating the fingertips [8,11,13] examined encoding of stimuli using abstract features of its response, such as mean firing rate, spike count or first-spike latency. Other models examined the evolution of synaptic weights between first- and second-order neurons over stimulus presentations but did not address the encoding of the stimuli [12]. In contrast, our models examine the encoding of tactile stimuli using synaptic integration, and thus provide mechanistic insight into how upstream neurons extract meaningful tactile information from the complex responses of the peripheral neuronal population.

We derive spiking models of first-order tactile neurons that fit and predict empirical data with good accuracy. We then simulate the first-order neuronal population innervating a fingertip, and investigate the computational capabilities afforded by synaptic integration of the population activity. We show that synaptic integration across first-order neurons can account for the human ability to discriminate tactile edge orientations with high acuity (< 3°) and speed [14]. Our results suggest that discriminating edge orientation in this manner relies on a small number of key inputs from first-order neurons. Our results also suggest that integrating fast-decaying (AMPA) synaptic input over a short time scale (i.e. coincidence detection) is critical for robustly discriminating with high acuity whereas integrating slow-decaying (NMDA) synaptic input serves to refine discrimination and maintain robustness over longer time scales (i.e. temporal summation).

## Results

### Data-driven models of first-order tactile neurons

We generated spiking models of human FA-1 neurons (n = 15 neurons) constrained by their response to oriented edges moving across the fingertip. Each model neuron innervated a set of mechanoreceptors, with axon spike initiation zone at each mechanoreceptor (Figure 1A). The input from each model mechanoreceptor depended on stimulus amplitude and distance (see Methods). Spikes were generated at each axon initiation zone following a linear relationship between mechanoreceptor input and axon spike rate, along with spike rate saturation (see Methods). The model neuron output followed a “reset” scheme, whereby spikes initiated at one initiation zone reset the other spike initiation zones (Figure 1A). The free parameters of the model neuron were the locations of the innervated mechanoreceptors (Figure 1B).

**Figure 1.**
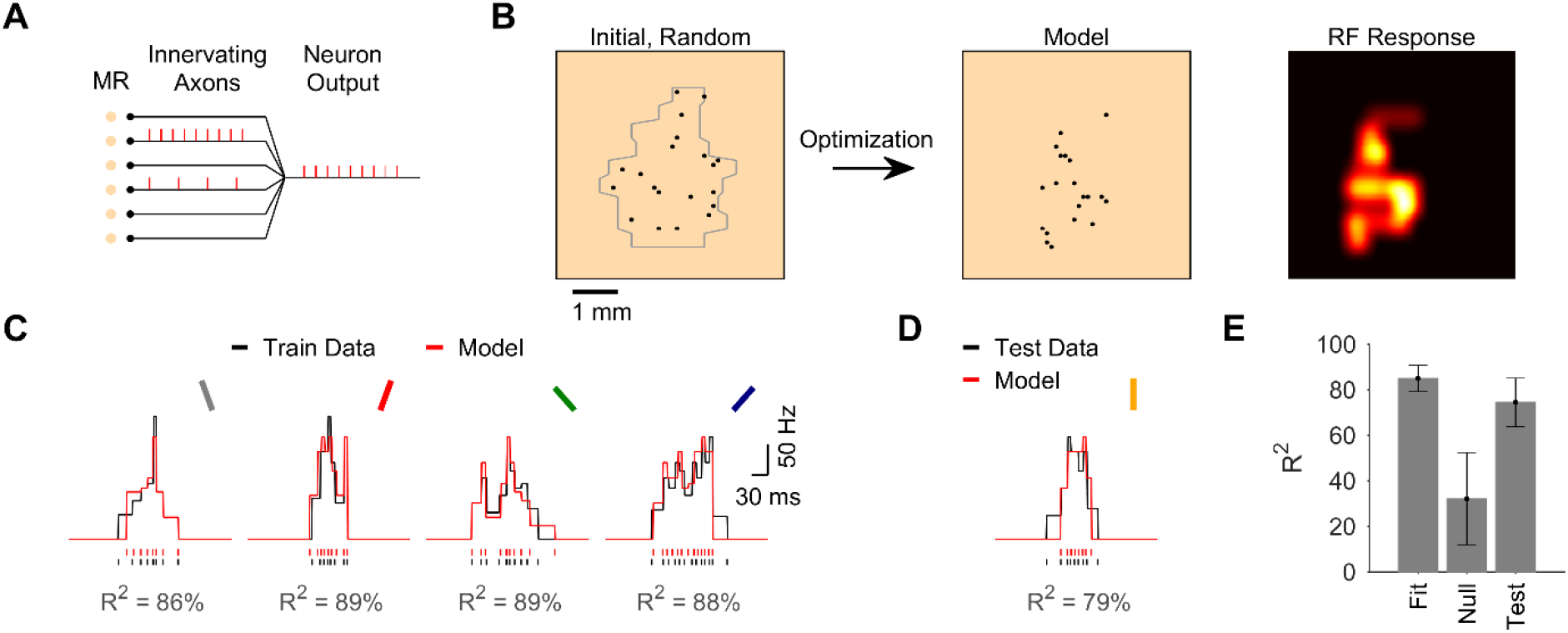
Data-driven models of first-order tactile neurons. **A.** The model neuron innervated a set of mechanoreceptors (MRs), each with its own axon. The model neuron output followed a “reset” scheme, whereby spikes triggered at one axonal spiking zone reset initiation at other spiking zones. **B.** The locations of innervated mechanoreceptors (black) were derived by the model optimization algorithm. At the start of the optimization, locations were random within the area derived from the recorded neuron’s response to a moving dot stimulus (gray, see Methods). The resulting model innervation pattern and receptive field (RF) response map are shown for the example FA-1 neuron whose response is illustrated in parts C and D. **C.** Fitness (R^2^) of model for example FA-1 neuron. Observed (black) and model (red) spike trains and rate curves in response to edges oriented −22.5, 22.5, −45, and 45°. **D.** Observed and model response to edge oriented 0°, which served to test the models. **E.** Model fitness, prediction accuracy, and the prediction accuracy of null models (shuffled, see Methods) across all neurons (n = 15). Prediction accuracy of models was significantly higher than that of null models (*p* < 10^-6^, paired-sample t-test). Error bars depict standard deviation.

We constrained the model for each recorded neuron using empirical spike responses to edges at four different orientations relative to their movement direction (±22.5, ±45°). We used genetic algorithm to derive the mechanoreceptor locations that yielded the best fit of model neuron response to the experimental response (see Methods). We optimized models using different number of mechanoreceptors (10 – 40) and then tested them on a different edge orientation (0°) that was not used during the model optimization. We selected the model that best fitted the training data and best predicted the test data, with fewest number of mechanoreceptors (see Methods). The model neurons fit the data with high accuracy (R^2^ = 85 ± 6%, mean ± SD, n = 15 neurons, Figure 1C,E), and predicted the test data similarly well (R^2^ = 74 ± 11%, Figure 1D,E).

Models for the different neurons required 20 ± 5 mechanoreceptors to reproduce the recorded responses. Prediction accuracy was significantly better than null models in which the mechanoreceptor locations were shuffled across the recorded neuron’s receptive field boundary (R^2^ = 32 ± 20%, n = 15, *p* < 10^-6^, paired-sample t-test, Figure 1E). Prediction accuracy was also significantly better than when simply using the experimental response of the nearest edge (−22.5 or 22.5°) to predict response to the test edge (0°, R^2^ = 51 ± 17%, n = 15, *p* < 10^-4^, paired-sample t-test). Our model neurons using spiking and resetting, in agreement with FA-1 physiology (see Methods), were thus able to fit and predict single trial spike response data with high accuracy, and therefore suggest that the diverse response of the neurons does not have to rely on analogue summation used in previous simpler models of human tactile receptive field responsivity [6].

### Encoding edge orientation via synaptic integration of first-order neuronal population activity

We next determined whether synaptic integration across the activity of FA-1 neuronal population can be used to discriminate edge orientations accurately. We simulated a model population of 330 FA-1 neurons innervating a 10×10 mm patch of skin, with model neuronal population generated as randomly-rotated variations of the 15 model neurons (see Methods). We simulated the population response to oriented edges passing over the skin patch, during a task of edge-orientation discrimination (Figure 2A). The task was to discriminate between two edges oriented at ±θ (where θ = 1, 3, 5, 10, or 20°). The response of the first-order population was then convolved with a postsynaptic potential (PSP) waveform and fed into a classifier with two units (tuned to θ and −θ) that performed synaptic integration across the neuronal population (see Methods).

**Figure 2.**
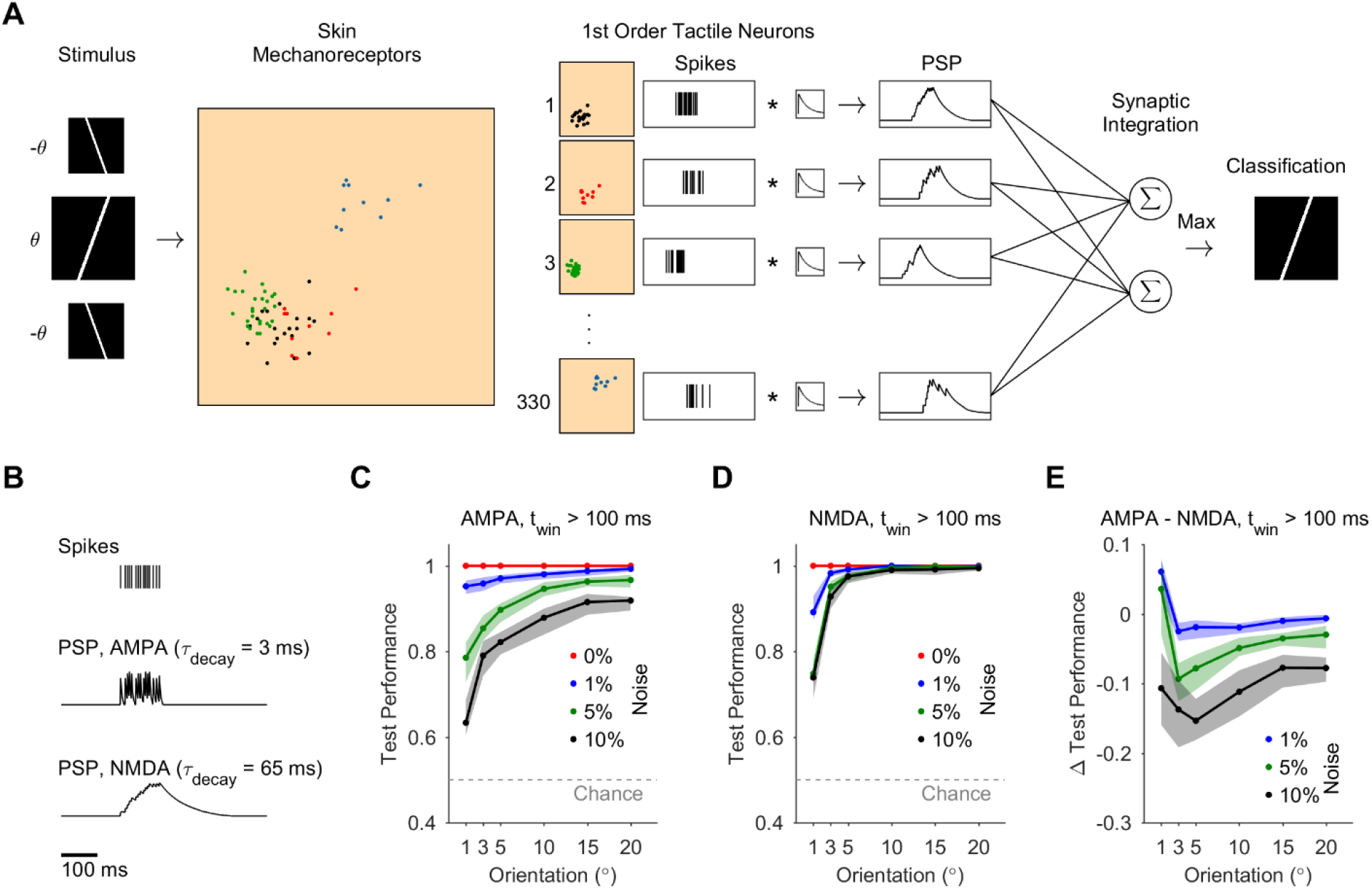
Discrimination of edge orientation using synaptic integration of first-order neuronal population activity. **A.** Simulated model population of FA-1 neurons innervating mechanoreceptors in a 10×10 mm patch of skin, during edge-orientation discrimination task (-θ vs θ, θ = 1-20°). Four example model first-order neurons are shown as color-coded sets of innervated mechanoreceptors, with their corresponding spike trains to the right. The neuronal population activity was fed into edge-orientation classifiers that performed synaptic integration of the neuronal population activity. The classifiers comprised of two units (tuned to −θ or θ) that performed a weighted sum of the synaptic inputs. Edge-orientation classification was determined according to the unit with maximal PSP value. **B.** PSPs were produced by convolving model first-order neuronal spike trains with a PSP waveform using either a short time constant (3 ms), corresponding to AMPA synapses, or a long time constant (65 ms), corresponding to NMDA synapses. **C, D.** Discrimination test performance using synaptic integration with AMPA (C) or NMDA (D) synapses. Each curve corresponds to a different level of stimulus noise (between 0 and 10%), and shows mean and 95% confidence intervals. Dashed gray line shows chance level (0.5). **E.** The difference in test performance between model networks using AMPA and NMDA synapses, for noise level 1, 5, 10%.

We inspected the discrimination accuracy of the classifiers using synaptic integration of PSPs of different decay time constants (see Methods) – either a short time constant corresponding to AMPA synapses (τ_decay_ = 3 ms), a long time constant corresponding to NMDA synapses (τ_decay_ = 65 ms), or a combination of both (Figure 2B). We also allowed negatively-weighted synapses with short and long time constants in the classifiers, which would correspond to feed-forward GABAA and GABAB inhibition, respectively. We inspected the discrimination accuracy under different levels of additive stimulus noise (0 – 10%, see Methods). We trained the classifiers using genetic algorithm to find the weights of the synaptic inputs from first-order neurons that yielded the best discrimination of edge orientations.

In the noiseless case, we found that synaptic integration of the activity in the first-order population could be used to discriminate different edge orientations perfectly with high acuity (to within ±1°), for either of the PSP time constants (Figure 2C,D). When adding noise to the stimulus, the discrimination accuracy of remained high (80-90% success), although discrimination became challenging for finer orientations. Discrimination accuracy using synaptic integration with NMDA synapses was more robust to noise compared to using AMPA synapses (improving test performance by 10 – 20%, Figure 2E). Synaptic integration using a combination of AMPA and NMDA PSPs did not significantly improve the robustness to noise.

We next varied the stimulus presentation time window (5, 10, 20, 50 ms, and unlimited, see Methods) and examined its effect on discrimination accuracy. In the noiseless case, the discrimination accuracy remained perfect regardless of the synapse type or time window size. When noise was applied, synaptic integration over shorter time windows increased the robustness of discrimination of classifiers used fast-decaying (AMPA) synapses, particularly for the fine angles (1 and 3°, Figure 3A). In contrast, integration time window did not significantly affect discrimination in classifiers that integrated slow-decaying (NMDA) synapses (Figure 3B). Overall, performance of classifiers that integrated population activity using AMPA synapses increased significantly in the case of short time window (5 ms, Figure 3D) compared to long time window (> 100ms, Figure 2C), approaching the robustness to noise of NMDA synapses over all orientations and improving performance for fine orientations (Figure 3E-F).

**Figure 3.**
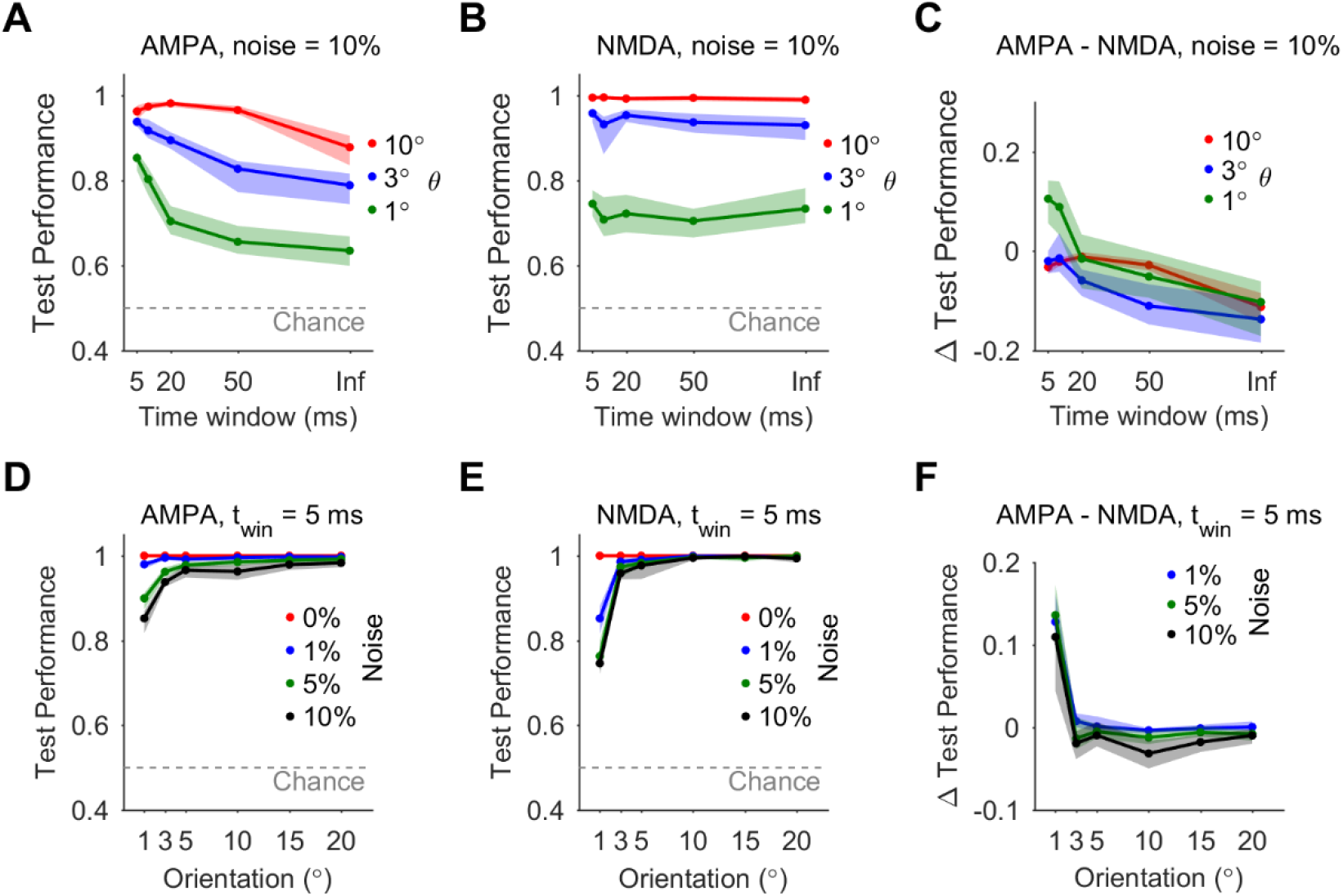
Integration of fast-decaying (AMPA) synapses over short time window robustly discriminates fine orientations. **A.** Discrimination test performance of classifiers integrating fast-decaying (AMPA) synaptic inputs under noise = 10%, as a function of stimulus presentation time window. **B.** Same as A but for classifiers integrating slow-decaying (NMDA) synaptic inputs. **C.** The difference in test performance between classifiers integrating AMPA synaptic inputs and classifiers integrating NMDA synaptic inputs. **D.** Discrimination test performance for classifiers integrating AMPA synaptic inputs over short integration time window (5 ms), for noise level = 0 – 10%. **E.** Same as D, but for classifiers integrating NMDA synaptic inputs. **F.** The difference in test performance between classifiers integrating AMPA synaptic inputs and classifiers integrating NMDA synaptic inputs, for integration time window of 5 ms and noise level 1, 5, 10%.

To inspect the contribution of first-order neurons to discrimination of different edges, we examined their synaptic weights and spatial layout in the different classifiers. Synaptic weights from most first-order neurons were close to 0 on average, and only a small subset of model first-order neurons consistently provided large contribution to classification (Figure 4A, based on 95% confidence intervals estimated by bootstrapping synaptic weights from 20 randomized trained classifiers, corrected for multiple comparisons). The number of key contributors was 11 ± 3 first-order neurons on average in classifiers using fast-decaying (AMPA) synapses, and similar (8 ± 2) in classifiers using slow-decaying (NMDA) synapses. We found no significant correlation between the edge orientation and the number of key synapses (*r* = 0.27, *p* = 0.6 for classifiers using AMPA synapses; *r* = 0.44, *p* = 0.38 for classifiers using NMDA synapses). The weights of the key excitatory synapses were anticorrelated between classifier units tuned to opposite orientations (*r* = −0.81 ± 0.09 across classifiers using AMPA synapses; *r* = −0.79 ± 0.03 across classifiers using NMDA synapses). An example of the key synaptic weights in classifiers integrating AMPA synaptic inputs for discriminating −20 and 20° is shown in Figure 4B (*r* = −0.96, *p* < 10^-4^). This was seen also in the receptive field map of the classifier units, which exhibited areas of positive and null sensitivity that alternated between oppositely-tuned units (Figure 4C). In addition, the receptive field map showed that the synaptic integration relied on coincidence of activation from distal parts of the edge stimulus, where the difference between opposite-oriented edges is largest.

**Figure 4.**
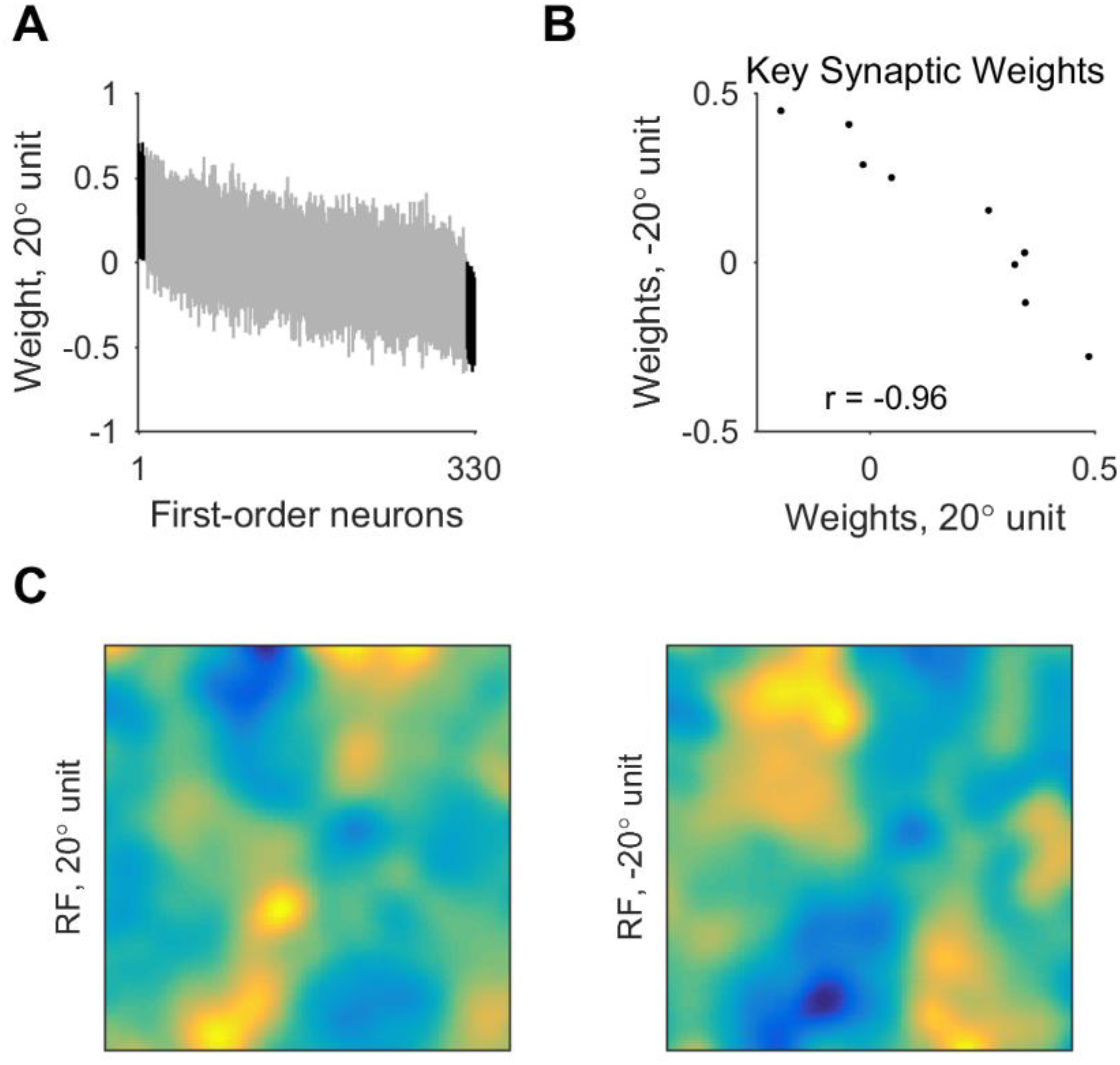
Synaptic weights and receptive field of edge-orientation classifiers. **A.** Synaptic weights from model first-order neurons onto classifier units using fast-decaying (AMPA) synapses and tuned to 20°, with presentation time window of 50 ms and noise level 5%. Shown are 95% confidence intervals of weights over 20 classifiers. Synaptic weights are shown in decreasing order of average weight over the 20 classifiers. Black – key synaptic weights, which were significantly different than 0. **B.** Key excitatory synaptic weights in classifiers using AMPA synapses and tuned to 20 vs −20°. **C.** Spatial layout of the synaptic weights in classifier units using AMPA synapses and tuned to −20 or 20°, with presentation time window of 50 ms and noise level 5%. Receptive fields represent averages over 20 classifiers.

## Discussion

Our study provides new insight into tactile processing at the level of the first-order neuronal population. We derived spiking models of human FA-1 neurons innervating mechanoreceptors in the fingertips that were able to fit and predict responses to oriented edge stimuli with high accuracy. We then simulated the first-order neuronal population activity from the fingertip and showed that its readout via synaptic integration could underlie human ability of using edge-orientation information with high acuity and speed, as in the context of hand control.

We investigated edge-orientation discrimination using synaptic integration as the readout of model first-order neuronal population activity, and thus were able to provide additional mechanistic insight and testable predictions unavailable to previous models that computed using abstract features of the population response, such as mean firing rate, spike count, or spike latency [8–11]. Our modeling approach predicts that the high accuracy and speed of edge-orientation discrimination could be largely mediated by synaptic integration of first-order tactile neuronal population activity, as early as the level of second-order tactile neurons at the spinal cord or cuneate nucleus [15]. Moreover, our modeling approach differentiated between integration mediated by fast-decaying (AMPA) synaptic inputs (leaning more towards coincidence detection), and slow-decaying (NMDA) synaptic inputs (leaning more towards summation).

Integration of AMPA-like synaptic inputs enabled robust discrimination especially of fine angles within a short time window. Therefore, AMPA inputs may mediate initial rapid computations based on bottom-up input, which could contribute to the fast responses that occur in the context of hand control [14]. Integration of NMDA-like synaptic inputs refined discrimination and maintained robustness when integrating inputs over longer time windows. Therefore, NMDA inputs may mediate the discrimination of objects based on their macro-geometric features – for example, discriminating between your house key and car key when searching in your pocket. The differentiation between AMPA and NMDA input streams could be established for example via compartmentalized synaptic locations on the postsynaptic neuron dendrites [16] or by multiplexed neural coding [17]. Our models also suggest a tendency for sparseness and the involvement of a small set of key synaptic inputs from first-order neurons, supporting previous empirical work [18] and suggesting that imposing a sparseness constraint on the synaptic weights may improve discrimination performance [19]. Previous studies also showed opposing areas of sensitivity in the receptive fields of cortical neurons [20]. Our models predict similar properties for second-order neurons that integrate first-order population activity, whereby key synaptic connections would be anticorrelated between oppositely-tuned neurons.

Our single-neuron models required ~20 mechanoreceptors on average to fit the spike response, which agrees with previous empirical estimates of the number of Meissner corpuscles converging onto a single FA-1 neuron [4]. We expect that a more complete set of constraints on the model neuron in the future would further increase the match between model and experimental estimates. Compared to the previous prediction accuracy of average spike response using convolution of the recorded receptive field responsivity [6], our model neurons enabled a high single-trial prediction accuracy and included physiological detail of mechanoreceptor innervation and spiking. An example of the added value of nonlinearity of spiking was that our models exhibited a variety of peak rates in response to different edge orientations, consistent with previous empirical results [6,7], thus indicating that the diverse response could be reproduced with a physiologically-realistic model neuron that used spiking and resetting [21–23], and did not depend on analogue summation used by previous simpler models of human tactile receptive field responsivity [6]. This richness arises because neuronal spike output is not only a function of the present input but also of the neuron’s previous spikes – a demonstration of how spiking increases a neuron’s computational power [24]. It remains to be seen if a spike reset scheme applies to other first-order tactile neurons, or in response to stimuli of lower intensity, which could instead involve a “mixing” of the spike trains [25].

Our models included skin mechanics only implicitly, and thus we kept the distance parameters of mechanoreceptor responsivity free in the model optimization. Although this simplification was sufficient for fitting and predicting spike responses in our data, our data-driven model optimization framework can be extended to include explicit components of skin dynamics as used in other tactile neuron models [10,23].

We have implemented additive noise at the level of the stimulus. The response of FA-1 and second-order cuneate neurons shows little variability [18,26,27], therefore there seems to be no significant noise at the level of mechanoreceptor to first-order neuron, or the synaptic projections from first-order neurons. While stimulus noise in the experimental data was < 1%, the larger levels of noise that we have investigated could represent other sources of noise such as variability in finger positioning during tasks. These in turn can account for the behavioral noise measured in tactile discrimination tasks [14].

## Methods

### Electrophysiology data

We used microneurography data of spike recordings from first-order tactile neurons, which was previously published [6]. The data consisted of 19 fast-adapting type 1 (FA-1) neurons from 13 human subjects. We utilized 15 of the 19 neurons, omitting two neurons whose receptive field was larger than the inter-stimulus spacing, and two neurons that had a low stimulus/non-stimulus response ratio. The recorded neurons innervated the glabrous skin of the index, long or ring finger. Neurons were stimulated via a rotating drum embossed with dots and edges at different orientations.

### First-order tactile neuron models

Model neurons innervated a subset of mechanoreceptors in a patch of skin modeled as a grid of mechanoreceptors located at 0.1 mm intervals [4]. The input from the model mechanoreceptor was proportional to the stimulus amplitude (indentation) and decreased with the distance from stimulus, following a sigmoidal function:

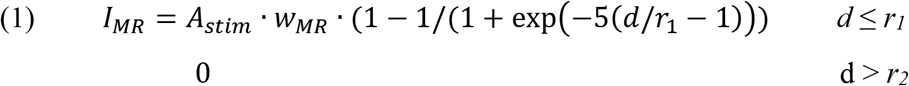

*A_stim_* was the stimulus amplitude, or tactile edge indentation, which was 0.5 mm as in the experimental data. *w_MR_* was the input weight of the mechanoreceptor, which was the same for all mechanoreceptors and set as twice the maximal firing rate of the experimental neuron. Thus, a stimulus indented 0.5 mm passing across a mechanoreceptor provided sufficient input to support the maximal firing rate of the recorded neuron. *d* was the distance (in mm) of the stimulus from the mechanoreceptor. *r_1_* was the sigmoidal half-height distance, and *r_2_* was the extent of the mechanoreceptor responsivity.

The model neuron innervated each mechanoreceptor with a dedicated axon [4] and spikes were initiated at each axon terminal (Figure 1A) [21–23,25,28,29]. Spike generation depended on mechanoreceptor input and the time from the last spike, following a linear relationship between mechanoreceptor input and spike rate (with gain = 1), and saturation at the maximal rate of the recorded neuron. The model neuron output followed a “reset” scheme, whereby spikes from one spiking zone propagated retrogradely to the other spiking zones and reset their spike initiation [21–23]. The model also included spike adaptation, whereby each model neuron axon fired only when input from mechanoreceptor was increasing, and was silent when the input remained the same or decreased. We used a spike threshold of 0.01 mm stimulus indentation [3]. We simulated the model neuron response to moving edge stimuli at Δt = 1 ms intervals. The model was implemented and simulated in Matlab (Mathworks).

### Neuron model optimization

We constrained the models with spike recordings of the response to edges of four different orientations (± 22.5, ± 45°). The error measure for the goodness of fit was R^2^ computed between the recorded and model spike rate curves:

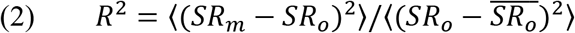

Where *SR_m_* is the spike rate time-series of the model neuron, and *SR_o_* is the spike rate time-series of the recorded neuron. The error was calculated for each edge orientation and averaged over the orientations to produce a ranking of the model.

We used a genetic algorithm [30–32] to search for the free model parameters, which were the locations of the mechanoreceptors innervated by the model neuron, as well as the two distance parameters of the mechanoreceptor responsivity (*r_1_* and *r_2_*, see above). The set of possible mechanoreceptor locations was delineated by the area of responsivity of the recorded neuron to a small dot stimulus (Figure 1B, gray). The search limits for the mechanoreceptor response distance parameters were [0.05,1] mm for *r_1_* and [0.2,1] mm for *r_2_*. During optimization, a population of 100 models was mutated with a probability of 0.1, and underwent crossover with a probability of 0.1 at each iteration. Models were selected based on how well they fitted the recorded response to the five edges (see above). We implemented the algorithm in Matlab, and optimization runtime of 500 iterations on 4 processors was 1 hour on average.

### Model neuron testing

After the optimization, we tested models using the spike recordings in response to edges oriented 0°. We compared the prediction to null models, in which the mechanoreceptor locations were shuffled across the area of responsivity (see above). The error measure was R^2^, as described above for the fitness. In addition, we compared the model prediction to that obtained by simply using the recorded response to either of the nearest edges ±22.5°.

### Model neuron selection

For each recorded neuron, we ran separate model optimizations using different number of innervated mechanoreceptors: 10, 20, 30, or 40. We selected from the resultant models the one that best fitted the training data and best predicted the test data, with fewest mechanoreceptors. We first selected the set of best models in terms of training data fitness, i.e. models with R^2^ within 5% of the maximal fit for each of the four edge orientations used to train the models. From this set, we selected the models that predicted test data with highest accuracy (R^2^ within 5% of the maximal prediction accuracy) and fewest mechanoreceptors (within 5 mechanoreceptors from minimum across the models). From these models, we selected the model with best prediction accuracy as the model for the recorded neuron.

### Model neuronal population

We generated model first-order neuronal population innervating a fingertip as randomly-rotated variations of the 15 model neurons. We tiled the model neurons over a 10×10 mm patch of skin, so that they innervated the patch at a uniform density of 140 neurons/cm^2^ [2]. This resulted in 330 model neurons innervating mechanoreceptors in the skin patch.

### Edge-orientation Discrimination

We simulated the activity of the neuronal population during a task of discriminating between two edges sweeping over the fingertip and oriented at −θ or θ, where θ = 1, 3, 5, 10, 15 or 20°. To discriminate the edge-orientations, we used classifiers comprised of two units, one tuned to −θ and another tuned to θ. Each unit performed a synaptic readout of neuronal population activity, via a weighted sum of the postsynaptic potentials (PSPs). In accordance with the physiology of first-order neuron projections, we used synapses with time constants that corresponded to AMPA and NMDA synapses [33,34]. The PSP vector from each first-order neuron was computed by convolving the spike train with PSP waveform, which was the difference of two exponentials:

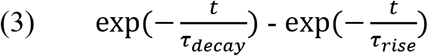

Where τ_rise_ = 0.5 ms, and τ_decay_ was either 3 ms as in AMPA receptors [35] or 65 ms as in NMDA receptors [36]. We also allowed fast-decaying and slow-decaying inhibitory synapses in the classifiers, which would correspond to feed-forward inhibition via GABAA and GABAB, respectively, with similar time constants to their excitatory counterparts [37,38]. Discrimination was based on the classifier unit with maximal PSP value over the time series, corresponding to a linear integration with threshold. We trained the model networks using a genetic algorithm to search for the input weights from the first-order neurons [39]. For the genetic algorithm, we used weights limits of [-1,1], a population of 100 models, mutation probability of 0.1, crossover probability of 0.1, and 200 iterations.

We investigated the discrimination accuracy under different levels of additive stimulus noise: 0, 1, 5, or 10%. Noise was added as 0.4 x 0.4 mm patches that varied in amplitude. For example, a noise level of 10% involved addition of noise patches of random amplitudes ranging between −10 and 10% of stimulus amplitude. While the additive noise could reduce stimulus indentation at some points, we forced the resulting stimulus to be strictly non-negative. We simulated 100 trials for each orientation, where trials differed in the noise added to the stimulus.

We picked 50 trials of each orientation at random for training the model networks (100 trials in total), and used the remaining 50 trials of each orientation for testing the model networks. For each task, we generated 20 model networks using different random subsets of training/testing trials. The performance measure was success discrimination rate over trials.

### Stimulus presentation time window

We investigated a range of time windows available for the edge-orientation discrimination. In the unlimited window case, edges passed over the skin patch fully from one end to the other. During shorter time windows, the edge passed for a limited time (5, 10, 20, or 50 ms) around the center of the skin patch.

### Statistical tests

We determined correlation using Pearson’s correlation coefficient. We tested for statistical significance using either paired or two-sample t-test, in Matlab. P-values < 0.05 were deemed significant. When estimating 95% confidence intervals, we used bootstrap estimation of the mean value in Matlab, and corrected for multiple comparisons using Bonferroni method (multiplying the *p* value by the number of comparisons).

## Acknowledgements

This work was supported by the Canadian Institutes of Health Research (Foundation Grant to JAP: 353197) and the Government of Ontario (Early Researcher Award to JAP). EH received a postdoctoral fellowship from the Brain and Mind Institute at Western University. JAP received a salary award from the Canada Research Chairs Program.

## References

1. Chemnitz A, Dahlin LB, Carlsson IK. Consequences and adaptation in daily life - patients’ experiences three decades after a nerve injury sustained in adolescence. BMC Musculoskelet Disord. 2013;14: 252. doi:10.1186/1471-2474-14-252

2. Johansson RS, Flanagan JR. Coding and use of tactile signals from the fingertips in object manipulation tasks. Nat Rev Neurosci. 2009;10: 345–359. doi:10.1038/nrn2621

3. Vallbo AB, Johansson RS. Properties of cutaneous mechanoreceptors in the human hand related to touch sensation. Hum Neurobiol. 1984;3: 3–14.

4. Nolano M, Provitera V, Crisci C, Stancanelli A, Wendelschafer-Crabb G, Kennedy WR, et al. Quantification of myelinated endings and mechanoreceptors in human digital skin. Ann Neurol. 2003;54: 197–205. doi:10.1002/ana.10615

5. Phillips JR, Johansson RS, Johnson KO. Responses of human mechanoreceptive afferents to embossed dot arrays scanned across fingerpad skin. J Neurosci Off J Soc Neurosci. 1992;12: 827–839.

6. Pruszynski JA, Johansson RS. Edge-orientation processing in first-order tactile neurons. Nat Neurosci. 2014;17: 1404–1409. doi:10.1038/nn.3804

7. Suresh AK, Saal HP, Bensmaia SJ. Edge orientation signals in tactile afferents of macaques. J Neurophysiol. 2016;116: 2647–2655. doi:10.1152/jn.00588.2016

8. Johansson RS, Birznieks I. First spikes in ensembles of human tactile afferents code complex spatial fingertip events. Nat Neurosci. 2004;7: 170–177. doi:10.1038/nn1177

9. Sripati AP, Bensmaia SJ, Johnson KO. A continuum mechanical model of mechanoreceptive afferent responses to indented spatial patterns. J Neurophysiol. 2006;95: 3852–3864. doi:10.1152/jn.01240.2005

10. Lesniak DR, Gerling GJ. Predicting SA-I mechanoreceptor spike times with a skin-neuron model. Math Biosci. 2009;220: 15–23. doi:10.1016/j.mbs.2009.03.007

11. Gerling GJ, Rivest II, Lesniak DR, Scanlon JR, Wan L. Validating a population model of tactile mechanotransduction of slowly adapting type I afferents at levels of skin mechanics, single-unit response and psychophysics. IEEE Trans Haptics. 2014;7: 216–228. doi:10.1109/TOH.2013.36

12. Rongala UB, Spanne A, Mazzoni A, Bengtsson F, Oddo CM, Jörntell H. Intracellular Dynamics in Cuneate Nucleus Neurons Support Self-Stabilizing Learning of Generalizable Tactile Representations. Front Cell Neurosci. 2018;12: 210. doi:10.3389/fncel.2018.00210

13. Saal HP, Delhaye BP, Rayhaun BC, Bensmaia SJ. Simulating tactile signals from the whole hand with millisecond precision. Proc Natl Acad Sci. 2017;114: E5693–E5702. doi:10.1073/pnas.1704856114

14. Pruszynski JA, Flanagan JR, Johansson RS. Fast and accurate edge orientation processing during object manipulation. eLife. 2018;7: e31200. doi:10.7554/eLife.31200

15. Jones EG. Cortical and Subcortical Contributions to Activity-Dependent Plasticity in Primate Somatosensory Cortex. Annu Rev Neurosci. 2000;23: 1–37. doi:10.1146/annurev.neuro.23.1.1

16. Larkum ME, Nevian T, Sandler M, Polsky A, Schiller J. Synaptic Integration in Tuft Dendrites of Layer 5 Pyramidal Neurons: A New Unifying Principle. Science. 2009;325: 756–760. doi:10.1126/science.1171958

17. Lankarany M, Al-Basha D, Ratté S, Prescott SA. Differentially synchronized spiking enables multiplexed neural coding. Proc Natl Acad Sci. 2019;116: 10097–10102. doi:10.1073/pnas.1812171116

18. Bengtsson F, Brasselet R, Johansson RS, Arleo A, Jörntell H. Integration of Sensory Quanta in Cuneate Nucleus Neurons In Vivo. PLOS ONE. 2013;8: e56630. doi:10.1371/journal.pone.0056630

19. Hay E, Ritter P, Lobaugh NJ, McIntosh AR. Multiregional integration in the brain during restingstate fMRI activity. PLoS Comput Biol. 2017;13: e1005410. doi:10.1371/journal.pcbi.1005410

20. DiCarlo JJ, Johnson KO, Hsiao SS. Structure of Receptive Fields in Area 3b of Primary Somatosensory Cortex in the Alert Monkey. J Neurosci. 1998;18: 2626–2645. doi:10.1523/JNEUROSCI.18-07-02626.1998

21. Horch KW, Whitehorn D, Burgess PR. Impulse generation in type I cutaneous mechanoreceptors. J Neurophysiol. 1974;37: 267–281. doi:10.1152/jn.1974.37.2.267

22. Fukami Y. Interaction of impulse activities originating from individual Golgi tendon organs innervated by branches of a single axon. J Physiol. 1980;298: 483–499.

23. Lesniak DR, Marshall KL, Wellnitz SA, Jenkins BA, Baba Y, Rasband MN, et al. Computation identifies structural features that govern neuronal firing properties in slowly adapting touch receptors. eLife. 2014;3: e01488. doi:10.7554/eLife.01488

24. Maass W. To Spike or Not to Spike: That Is the Question. Proc IEEE. 2015;103: 2219–2224. doi:10.1109/JPROC.2015.2496679

25. Goldfinger MD, Fukami Y. Interaction of activity in frog skin touch afferent units. J Neurophysiol. 1981;45: 1096–1108. doi:10.1152/jn.1981.45.6.1096

26. Jörntell H, Bengtsson F, Geborek P, Spanne A, Terekhov AV, Hayward V. Segregation of Tactile Input Features in Neurons of the Cuneate Nucleus. Neuron. 2014;83: 1444–1452. doi:10.1016/j.neuron.2014.07.038

27. Hayward V, Terekhov AV, Wong S-C, Geborek P, Bengtsson F, Jörntell H. Spatio-temporal skin strain distributions evoke low variability spike responses in cuneate neurons. J R Soc Interface. 2014;11: 20131015. doi:10.1098/rsif.2013.1015

28. Eagles JP, Purple RL. Afferent fibers with multiple encoding sites. Brain Res. 1974;77: 187–193.

29. Lindblom Y, Tapper DN. Integration of impulse activity in a peripheral sensory unit. Exp Neurol. 1966;15: 63–69.

30. Druckmann S, Banitt Y, Gidon A, Schürmann F, Markram H, Segev I. A Novel Multiple Objective Optimization Framework for Constraining Conductance-Based Neuron Models by Experimental Data. Front Neurosci. 2007;1: 7–18. doi:10.3389/neuro.01.1.1.001.2007

31. Hay E, Hill S, Schürmann F, Markram H, Segev I. Models of neocortical layer 5b pyramidal cells capturing a wide range of dendritic and perisomatic active properties. PLoS Comput Biol. 2011;7: e1002107. doi:10.1371/journal.pcbi.1002107

32. Hay E, Schürmann F, Markram H, Segev I. Preserving axosomatic spiking features despite diverse dendritic morphology. J Neurophysiol. 2013;109: 2972–2981. doi:10.1152/jn.00048.2013

33. Soto C, Aguilar J, Martín-Cora F, Rivadulla C, Canedo A. Intracuneate mechanisms underlying primary afferent cutaneous processing in anaesthetized cats. Eur J Neurosci. 2004;19: 3006–3016. doi:10.1111/j.0953-816X.2004.03432.x

34. Kus L, Saxon D, Beitz AJ. NMDA R1 mRNA distribution in motor and thalamic-projecting sensory neurons in the rat spinal cord and brain stem. Neurosci Lett. 1995;196: 201–204.

35. Hestrin S. Activation and desensitization of glutamate-activated channels mediating fast excitatory synaptic currents in the visual cortex. Neuron. 1992;9: 991–999.

36. Rhodes P. The properties and implications of NMDA spikes in neocortical pyramidal cells. J Neurosci Off J Soc Neurosci. 2006;26: 6704–6715. doi:10.1523/JNEUROSCI.3791-05.2006

37. Salin PA, Prince DA. Electrophysiological mapping of GABAA receptor-mediated inhibition in adult rat somatosensory cortex. J Neurophysiol. 1996;75: 1589–1600. doi:10.1152/jn.1996.75.4.1589

38. Gerrard LB, Tantirigama MLS, Bekkers JM. Pre- and Postsynaptic Activation of GABAB Receptors Modulates Principal Cell Excitation in the Piriform Cortex. Front Cell Neurosci. 2018;12. doi:10.3389/fncel.2018.00028

39. Such FP, Madhavan V, Conti E, Lehman J, Stanley KO, Clune J. Deep Neuroevolution: Genetic Algorithms Are a Competitive Alternative for Training Deep Neural Networks for Reinforcement Learning. ArXiv171206567 Cs. 2017; Available: http://arxiv.org/abs/1712.06567

